# ATG9 vesicles comprise the seed membrane of mammalian autophagosomes

**DOI:** 10.1101/2022.08.16.504143

**Authors:** Taryn J. Olivas, Yumei Wu, Shenliang Yu, Lin Luan, Peter Choi, Shanta Nag, Pietro De Camilli, Kallol Gupta, Thomas J. Melia

## Abstract

During autophagosome biogenesis, the incorporation of transmembrane proteins into the expanding phagophore is not readily observed. In addition, the membrane surface area of the organelle expands rapidly, while the volume of the autophagosome is kept low. Several recent studies have suggested a model of membrane expansion that explains how these attributes are maintained. The autophagosome expands predominantly through the direct protein-mediated transfer of lipids through the lipid transfer protein ATG2. As these lipids are only introduced into the cytoplasmic-facing leaflet of the expanding phagophore, full membrane growth also requires lipid scramblase activity. ATG9 has been demonstrated to harbor scramblase activity and is essential to autophagosome formation, however if and when it is integrated into mammalian autophagosomes remains unclear. Here we show that in the absence of lipid transport, ATG9 vesicles are already fully competent to collect proteins normally found on mature autophagosomes, including LC3-II. Further, through the novel use of styrene-maleic acid lipid particles as a nanoscale interrogation of protein organization on intact membranes, we show that ATG9 is fully integrated in the same membranes as LC3-II, even on maturing autophagosomes. The ratios of these two proteins at different stages of maturation demonstrate that ATG9 proteins are not continuously integrated, but rather are present on the seed vesicles only and become diluted in the rapidly expanding autophagosome membrane. Thus, ATG9 vesicles are the seed membrane from which mammalian autophagosomes form.

## Introduction

Macroautophagy, herein referred to as autophagy, is the process of clearing large cytoplasmic debris such as protein aggregates and even whole dysfunctional organelles in order to maintain cellular homeostasis. This is mediated by the autophagosome— an organelle characterized by its *de novo* formation around targeted cytosolic cargoes, encapsulating and subsequently trafficking them to the lysosome for degradation. As autophagosomes are largely separate from classic vesicle trafficking pathways, how these organelles form *de novo* has been an area of intense exploration. Intriguingly, recent studies implicate an entirely novel method of membrane expansion: the direct delivery of lipid from a donor membrane to the autophagosome via protein-mediated lipid transport through the autophagy protein ATG2 (Maeda et al., 2019; Osawa et al., 2019; Valverde et al., 2019).

ATG2 is among a new class of bulk lipid transport proteins characterized by an N-terminal chorein domain and a long hydrophobic channel that makes it capable of transporting tens of lipids at once (Leonzino et al., 2021). This higher capacity to move large quantities of lipids suggests that bulk lipid flow may be suitable to grow an organelle, however lipid transfer alone cannot complete the process. In order for the membrane to expand, there must also be a mechanism to populate the inner leaflet of the bilayer with lipids. The presence of a lipid floppase or scramblase, proteins capable of moving lipids across the bilayer from the outer to inner leaflet, would fulfill this requirement. Recently, ATG9, the only transmembrane protein in the core autophagy machinery, was defined to function as a scramblase (Ghanbarpour et al., 2021; Maeda et al., 2020; Matoba et al., 2020), and to bind ATG2 proteins directly to support autophagosome expansion (Ghanbarpour et al., 2021; Gómez-Sánchez et al., 2018; Kotani et al., 2018; Tang et al., 2019). This makes ATG9 an excellent candidate to be the phagophore scramblase. However, it remains unclear whether ATG9 is ever resident on the mammalian autophagosome.

Recent structural studies identified a hydrophilic channel in ATG9 (Guardia et al., 2020; Maeda et al., 2020; Matoba et al., 2020), which in vitro assays on liposomes confirmed supported lipid scrambling activity (Ghanbarpour et al., 2021; Maeda et al., 2020; Matoba et al., 2020). Isolated Atg9 vesicles from yeast can also support the lipid-conjugation of the autophagy protein Atg8 *in vitro*, when all the conjugation machinery is supplied exogenously, indicating the surface of these vesicles is autophagy-competent (Sawa-Makarska et al., 2020). Critically, ATG9 must be incorporated into the membrane of the growing autophagosome to support the membrane expansion model. Biochemically-isolated autophagosomes from yeast appear to have a small number of Atg9 proteins (Yamamoto et al., 2012), but in mammals evidence for ATG9 on the autophagosome has been lacking. ATG9 is found on vesicles throughout the endolysosomal pathway (Imai et al., 2016; Puri et al., 2013; Zhou et al., 2022), as well as apposed to several membrane compartments including lipid droplets (Mailler et al., 2021), and even transit through the plasma membrane (Claude-Taupin et al., 2021). ATG9 vesicles also travel to and shuttle around phagophores when autophagy is initiated (Mari et al., 2010; Orsi et al., 2012; Young et al., 2006) but these vesicles appear to remain proximal to rather than integrated in growing phagophores (Orsi et al., 2012). Indeed, detection of any transmembrane proteins on the completed autophagosome is challenging (Fengsrud et al., 2000). Intriguingly, a consequence of driving membrane expansion predominantly through protein-mediated lipid transport rather than vesicle fusion is that transmembrane proteins would be largely excluded from the final mature organelle. Thus, ATG9 proteins might never accumulate at the autophagosome beyond whatever small number of proteins are present before lipid transport is activated. If ATG9-related vesicles comprised a seed membrane for autophagosome biogenesis, then the density of ATG9 proteins would be highest in the moments just before lipid transport occurs.

We previously generated cells with both forms of human ATG2 (ATG2A and ATG2B) knocked out (Valverde et al., 2019), which we speculate would accumulate the precursor to the phagophore membrane. Here we show that these cells harbor a pre-ATG2 compartment that is significantly enriched for ATG9 and LC3B-II. We then employ styrene maleic acid copolymer nanodiscs to capture small spans of membrane that biochemically show that ATG9 and LC3 are on the same membrane in both this pre-ATG2 compartment and on wild type autophagosomes. Together, these data establish that ATG9 is resident on the phagophore membrane, positioning it to receive lipids from ATG2 and populate the inner leaflet with lipid during autophagosome expansion (Figure 1A).

**Figure 1.**
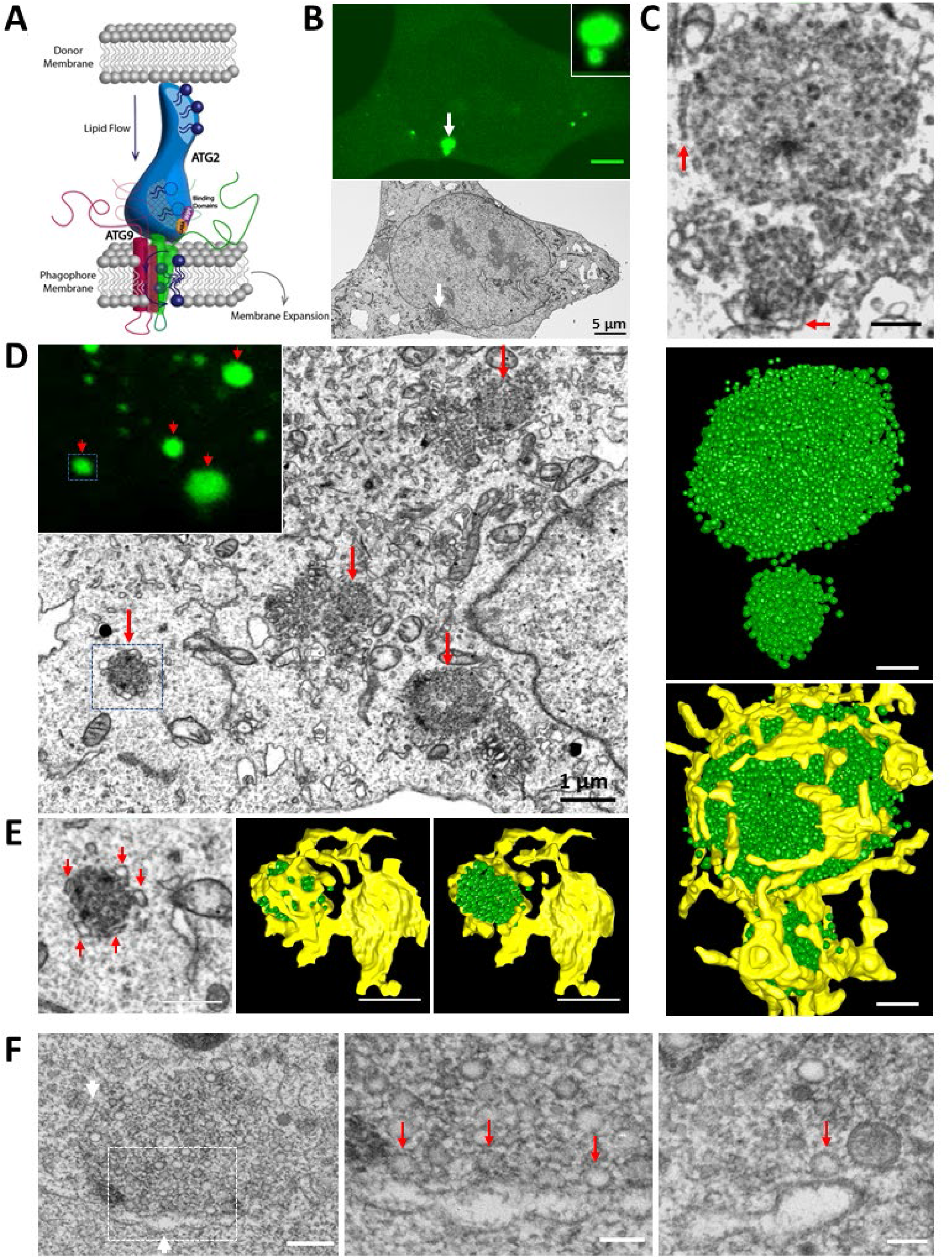
LC3B-positive vesicles accumulate in large clusters in ATG2 DKO cells. **(A)** Model for autophagosome growth via direct lipid transport by ATG2. ATG2 moves lipids from a donor membrane to the phagophore membrane, where ATG9 scrambles the lipids to populate both leaflets of the growing bilayer. This model requires that ATG9 be integrated in the expanding phagophore. **(B)** Correlative light electron microscopy (CLEM) combined with focused ion beam scanning electron microscopy (FIB-SEM) in ATG2 DKO HEK293 cells revealed two accumulations of GFP-LC3B signal (white arrow and inset) that correlated to the site of accumulated vesicles (white arrow on the cell of FIB-SEM image). Scale bar: 5 μm. **(C)** 3D segmentation of two vesicle clusters. Top: two vesicle clusters from FIB-SEM (B) at high magnification showing ER structures (red arrows) surrounding the vesicle clusters. Middle: segmented small vesicle clusters. Bottom: vesicle clusters with abundant ER contacts (yellow). Scale bar: 250 nm. **(D)** More FIB-SEM-imaged small vesicle clusters (red arrows) are correlated to the GFP-LC3B positive signals (inset, red arrows). Scale bar: 1 μm. **(E)** 3D segmentation of the vesicle accumulation from the blue box in (D). Left: abundant ER structures are closely surrounding the vesicle cluster (red arrows) on the FIB-SEM image. Middle and Right: 3D models are showing the vesicle cluster encaged by the ER structure. Scale bars: 500 nm. **(F)** Transmission electron microscopy (TEM) images showing the contacts between the small vesicles and ER. Left: one vesicle cluster with ER around (white arrows). Middle: the high magnification image from the Left inset showing the vesicles contacting ER without clear distance (red arrows). Right: a vesicle-ER contact with a space of 8 nm. Scale bar: 250 nm in Left; 100 nm in Middle and Right.

## Results

### Large vesicle clusters full of early autophagy proteins accumulate in mammalian ATG2 depleted cells

ATG2-mediated lipid transport is essential to autophagosome growth and suggests a simple model where ATG2 delivers lipids in bulk to a pre-existing progenitor or seed membrane (Figure 1A). To look for these progenitors, we removed ATG2 from the cell and examined what structures accumulate at the resulting sites of frustrated autophagosome biogenesis. Gene-edited HEK293 cells lacking both mammalian ATG2 genes (ATG2 double knockout or ATG2 DKO) accumulate abnormally large LC3B-positive immunofluorescent puncta but no mature autophagosomes (Valverde et al., 2019). To determine the ultrastructure of these LC3B-positive puncta, we employed correlative light electron microscopy (CLEM) combined with focused ion beam scanning electron microscopy (FIB-SEM) on ATG2 DKO cells expressing GFP-LC3B. In these cells, the GFP-LC3B puncta correlated with large, roughly spherical vesicle clusters (Figure 1, B and C; Movie S1). Segmentation and 3D reconstruction of these clusters indicated as many as several hundred vesicles accumulate at each site, and each cluster is in close contact with significant stretches of endoplasmic reticulum (ER) such that in some cases the ER appeared to partially encapsulate a cluster (Figure 1, C-E). Transmission electron microscopy revealed the vesicles at the edge of the cluster and the surrounding ER are separated by less than 10 nm (Figure 1F).

The LC3B at these sites is almost certainly in the lipidated form, as we detected virtually no LC3B-I in these cells (Valverde et al., 2019), thus at least some of these vesicles are LC3B-II-positive. We next used fluorescence microscopy to probe the for other early autophagy factors required for initiation of autophagosome formation. We detected ATG9A, p62, WIPI2 and FIP200 (Figure 2, A-H) colocalizing with the LC3B-positive puncta, consistent with previous reports indicating most autophagosome biogenesis proteins collect in large puncta when ATG2 is depleted (Bozic et al., 2020; Stanga et al., 2019; Tamura et al., 2017; Tang et al., 2017; Tang et al., 2019; Valverde et al., 2019; Velikkakath et al., 2012). As a transmembrane protein, ATG9A must be embedded within one or more vesicle population found here. ATG9A and p62 each appeared to be broadly colocalized with LC3B (Figure 2, A-D), though the distribution of each marker is not always homogeneous (Figure S1), and thus we cannot conclude whether ATG9A and LC3B-II decorate the same or different membranes. WIPI2, a PI3P-binding protein implicated in tethering phagophores to the ER (Dooley et al., 2014; Dooley et al., 2015; Polson et al., 2010; Zhao et al., 2017), encircled the LC3B signal (Figure 2, E and F; Figure S1 C), consistent with the organization of the ER in close contact at the periphery of the vesicle cluster. The autophagic scaffolding protein FIP200 was present in a less discernible pattern (Figure 2, G and H; Figure S1 D), but nonetheless indicates that initiation factors for autophagosome formation arrive at this site. We refer to these sites as pre-ATG2 compartments. Interestingly, autophagosome biogenesis at omegasomes is believed to involve the close apposition of the ER with expanding phagophores (Nascimbeni et al., 2017; Polson et al., 2010; Uemura et al., 2014). Thus here, the organization of ER tightly surrounding membranes laden with autophagy factors is broadly consistent with the structure of early autophagosome biogenesis sites (Uemura et al., 2014), and critically, the formation of lipid-anchored LC3B suggests a biochemically competent seed vesicle exists in this space.

**Figure 2.**
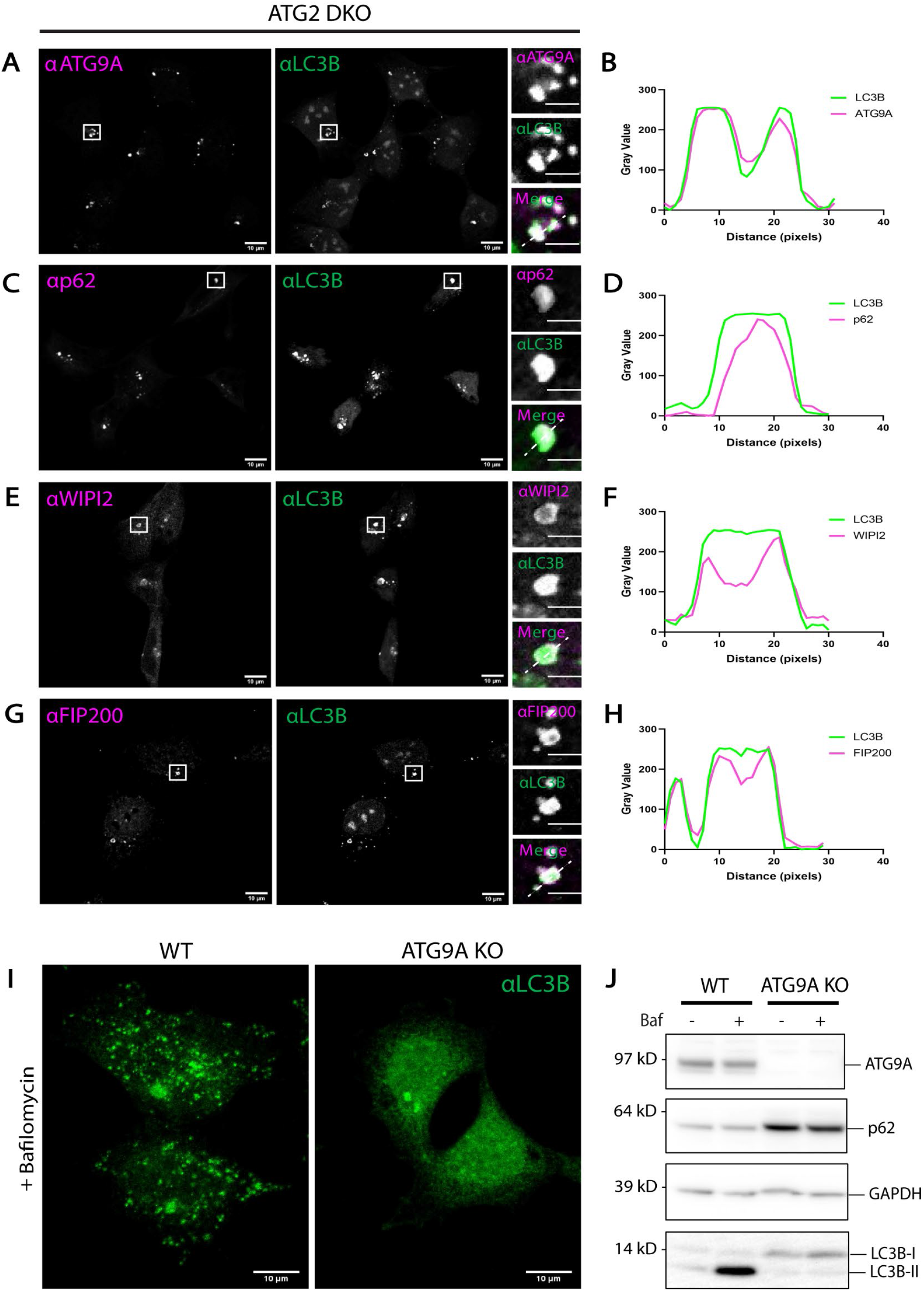
Early autophagy factors collect in the pre-ATG2 compartment. Immunofluorescence of LC3B and indicated early autophagy proteins in ATG2 DKO HEK293 cells. **(A-B)** ATG9A signal fills the compartment covering the span of the vesicle cluster suggesting ATG9 vesicles are present. **(C-D)** p62 signal is strongest in the center of the compartment but colocalizes with LC3B signal throughout. **(E-F)** WIPI2 signal surrounds the compartment, colocalizing with LC3B at the edges. **(G-H)** FIP200 signal is throughout the compartment suggesting autophagosome formation was initiated and halted before expansion with the loss of ATG2. All line scans correspond to the white dashed line in the insets. Scale bars: 10 μm. Inset scale bars: 3 μm. **(I)** Immunofluorescence of endogenous LC3B in WT and ATG9A KO HEK293 cells. The ATG9A KO does not show Bafilomycin A1-dependent accumulation of autophagosomes as in the WT, suggesting autophagy activity is impaired in ATG9A KO cells. Scale bars: 10 μm. **(J)** Immunoblot showing the loss of ATG9A protein in the ATG9A KO. LC3B lipidation is blocked and p62 accumulates, suggesting no autophagic flux. Loaded protein: 10 μg.

### Knock out of ATG9A results in little to no LC3B lipidation

Like ATG2, ATG9 is essential for autophagosome biogenesis (He et al., 2008; Orsi et al., 2012; Yamamoto et al., 2012). To determine where in the autophagosome biogenesis pathway ATG9A is needed, we used gene-editing to remove ATG9A in HEK293 cells (ATG9A KO). ATG9A KO cells did not form LC3B-positive punctate structures even with bafilomycin A1 treatment to accumulate autophagosomes (Figure 2I), and unlike ATG2 DKO cells, they did not accumulate LC3B-II. They also did not support autophagic flux; these cells had an accumulation of p62 and did not accumulate LC3B-II under bafilomycin A1 treatment (Figure 2J). These phenotypes can be partially rescued by transient expression of exogenous FLAG-tagged ATG9A (Figure S2, A and B), and stable expression of FLAG-ATG9A fully restored bafilomycin A1-dependent LC3B-II accumulation (Figure S2 B). Thus, in these cells, loss of ATG9A correlated with a loss of autophagy-related LC3B-II formation.

### ATG9A and LC3B reside in the same membrane fraction after density gradient fractionation

If ATG9 vesicles are integrated into developing autophagosomes at any point, ATG9 and LC3B-II will be found on the same membranes. If so, these markers should co-enrich during membrane fractionation. We used a well-established density gradient membrane fractionation protocol (Strømhaug et al., 1998) to isolate LC3B-positive membranes from wild type (WT) HEK293 cells that were also stably expressing GFP-LC3 to facilitate subsequent immunoisolation studies.

This technique involves several centrifugation steps to separate membrane compartments from one another (Figure 3A). Consistent with Strømhaug et al and our own previous results (Jeong et al., 2009; Nguyen et al., 2020; Strømhaug et al., 1998), GFP-LC3B-II and LC3B-II were strongly enriched in fraction 6 in WT cells (Figure 3, B, D, and G), hereafter referred to as the autophagosome (AP) fraction. Strikingly, ATG9A was also robustly enriched in this fraction (Figure 3, B, D and E). As autophagosomes are relatively scarce and not protein dense, the total protein recovered in the AP fraction is quite low. As such, even as ATG9A was enriched in this fraction, the vast majority of ATG9A was actually found in fractions 3-5, where other transmembrane proteins such as TOM20 and Calnexin were enriched (Figure 3, B and D) consistent with the predominantly non-autophagosomal distribution of ATG9A described by microscopy. Thus, biochemical enrichment revealed that a small fraction of the ATG9A pool strongly co-enriches with the autophagosome membrane population.

**Figure 3.**
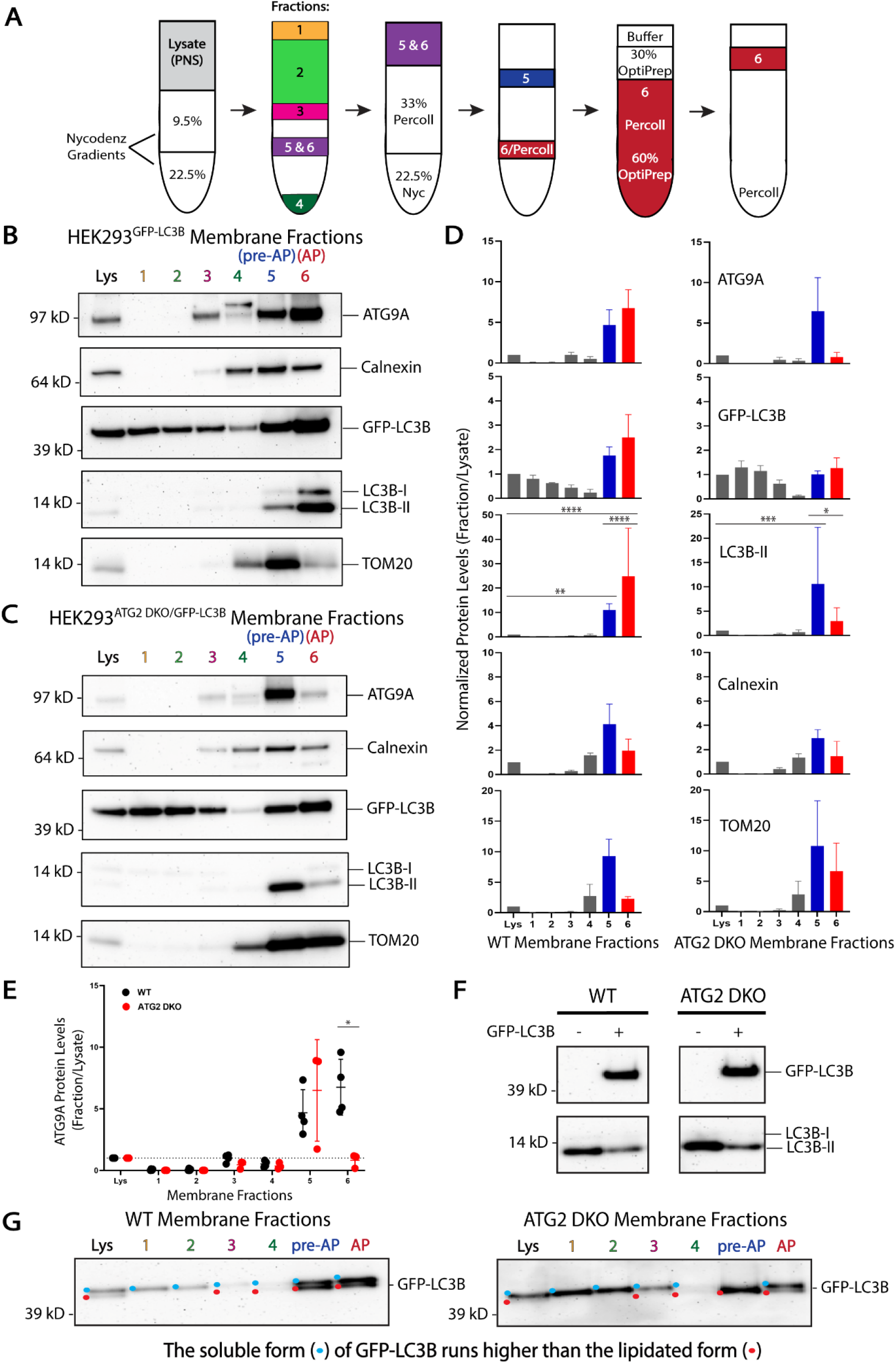
ATG9A and LC3B reside in the same membrane fractions enriched for autophagosomes or their precursor membranes. **(A)** Experimental scheme for density gradient membrane fractionation protocol (see Methods and (Strømhaug et al., 1998)). **(B)** Immunoblot of cell membrane fractions from WT HEK293 cells showing enrichment of ATG9A and LC3B together in fractions 5 and 6, fraction 6 being reportedly enriched for autophagosome membranes. Cells were grown in DMEM before being starved in EBSS and treated with 0.1 μM Bafilomycin A1 for 4-6 h. Loaded protein: 3 μg. **(C)** Immunoblot of cell membrane fractions from ATG2 DKO HEK293 cells showing enrichment of ATG9A and LC3B together in fraction 5, and pointedly fraction 6 is largely depleted of both proteins in this cell line blocking autophagosome formation. Cells were grown in DMEM and were not treated before collection (ATG2 DKO cells accumulate early autophagic factors without starvation or bafilomycin A1 treatment (Valverde et al., 2019)). Loaded protein: 3 μg. **(D)** Densitometric quantification of ATG9A, GFP-LC3B, LC3B-II, Calnexin, and TOM20 in each cell line membrane fractionation. The intensity of the bands in **B and C** were normalized to the lysate, and statistical significance was assessed by two-way ANOVA. *, adjusted p-value < 0.05. **, adjusted p-value < 0.01. ***, adjusted p-value < 0.001. ****, adjusted p-value < 0.0001. **(E)** Densitometric quantification of ATG9A in each fraction compared between the two cell lines. The intensity of the bands in **B** and **C** were normalized to the lysate, and statistical significance was assessed by multiple unpaired t-tests. *, adjusted p-value < 0.05. More quantifications are shown in **Figure S3**. **(F)** Immunoblot comparing endogenous levels of LC3B-II in HEK293 cells with or without overexpression of GFP-LC3B. The endogenous LC3B pool in ATG2 DKO cells is mostly but not completely in form II, perhaps because the ability to lipidate LC3B is saturated with GFP-LC3B overexpression. WT and ATG2 DKO samples are from the same blot. Loaded protein: 10 μg. **(G)** Immunoblots from membrane fractions from both WT and ATG2 DKO HEK293 cells were run on the Bolt SDS-PAGE system, which allows larger fusion proteins to run far enough to separate protein conjugated to lipid, to distinguish GFP-LC3B-I and GFP-LC3B-II. Colored dots indicate whether the non-lipidated (blue) and/or lipidated (red) forms of GFP-LC3B were detectable in each lane. Loaded protein: 3 μg.

In ATG2 DKO cells, most of the LC3B is form II ((Valverde et al., 2019) and Figure 3, F and G) and it colocalized with ATG9A in the pre-ATG2 compartment (Figure 2). If LC3B is directly lipidated on ATG9A vesicles in this compartment, that would indicate ATG9 vesicles are fully competent to promote the downstream biochemistry involved in the lipidation reaction as would be expected of a seed membrane. To study whether ATG9A and LC3B are on biochemically separable membranes, we again utilized stable expression of GFP-LC3B in ATG2 DKO cells. In ATG2 DKO cells, where autophagosome formation is largely disrupted, neither LC3B-II nor ATG9A enriched in the AP fraction (Figure 3, C, D, and E). Instead, virtually all of the ATG9A, LC3B-II and GFP-LC3B-II was recovered in fraction 5, (Figure 3, C, D, and E; Figure S3), which was also enriched for ER and mitochondrial markers. This fraction was originally described by Strømhaug et al as an “ER” fraction– in our hands we observed significant accumulation of autophagy markers in both the WT and ATG2 DKO samples (Figure 3, B-D) consistent with the presence of omegasomes or phagophores expected to be proximal to the ER, and thus we consider this to also be a pre-autophagosome-enriched (pre-AP) fraction.

Thus, ATG9A and LC3B were each resident in membranes of comparable density and further, when autophagosome biogenesis was specifically disrupted by the depletion of ATG2, they each redistributed to the same new density, strongly suggesting that in WT cells they were present on the same structure. However, the limitations of density gradient isolation are also obvious as both Calnexin and TOM20 can be readily detected in each LC3B-II fraction, necessitating a “higher resolution” biochemistry approach.

### Nanodiscs reveal that ATG9A and LC3B-II are on the same vesicle membrane in the pre-ATG2 compartment

Styrene maleic acid (SMA) copolymer is an anionic hypercoiling polymer that has drawn attention for its ability to stabilize small, intact regions of membrane for downstream analysis (Dörr et al., 2016). In an aqueous environment, the ring structure of the copolymer collapses exposing hydrophobic styrenes such that the polymer behaves as an amphipathic molecule (Tonge and Tighe, 2001). The styrene intercalates between the bilayer acyl chains similarly to cholesterol, destabilizing the membrane and capturing lipids in nanodiscs approximately 10 nm in diameter (Jamshad et al., 2015). Importantly, this differs dramatically from other membrane encapsulating nanodiscs because at no time does SMA extraction require detergent, therefore maintaining the integrity of the isolated membrane.

SMA is most commonly used as a preparative tool for the study of membrane-embedded proteins from native sources during structure analysis (Dörr et al., 2016; Esmaili and Overduin, 2018; Postis et al., 2015). Under those working conditions, membranes are fully solubilized into nanodiscs called SMA lipid particles (SMALPs). With artificial liposomes as a substrate, we observed this complete solubilization at 2.5% (w/v) SMA (Figure 4A). At lower concentrations of SMA, SMALPs of comparable dimensions still form (e.g. particles in 0.05% SMA), but largely intact membranes also persist. This demonstrates the utility of SMA to “punch out” single nanodomains from otherwise intact membrane, each estimated to comprise about 140 lipids on average (Jamshad et al., 2015).

**Figure 4.**
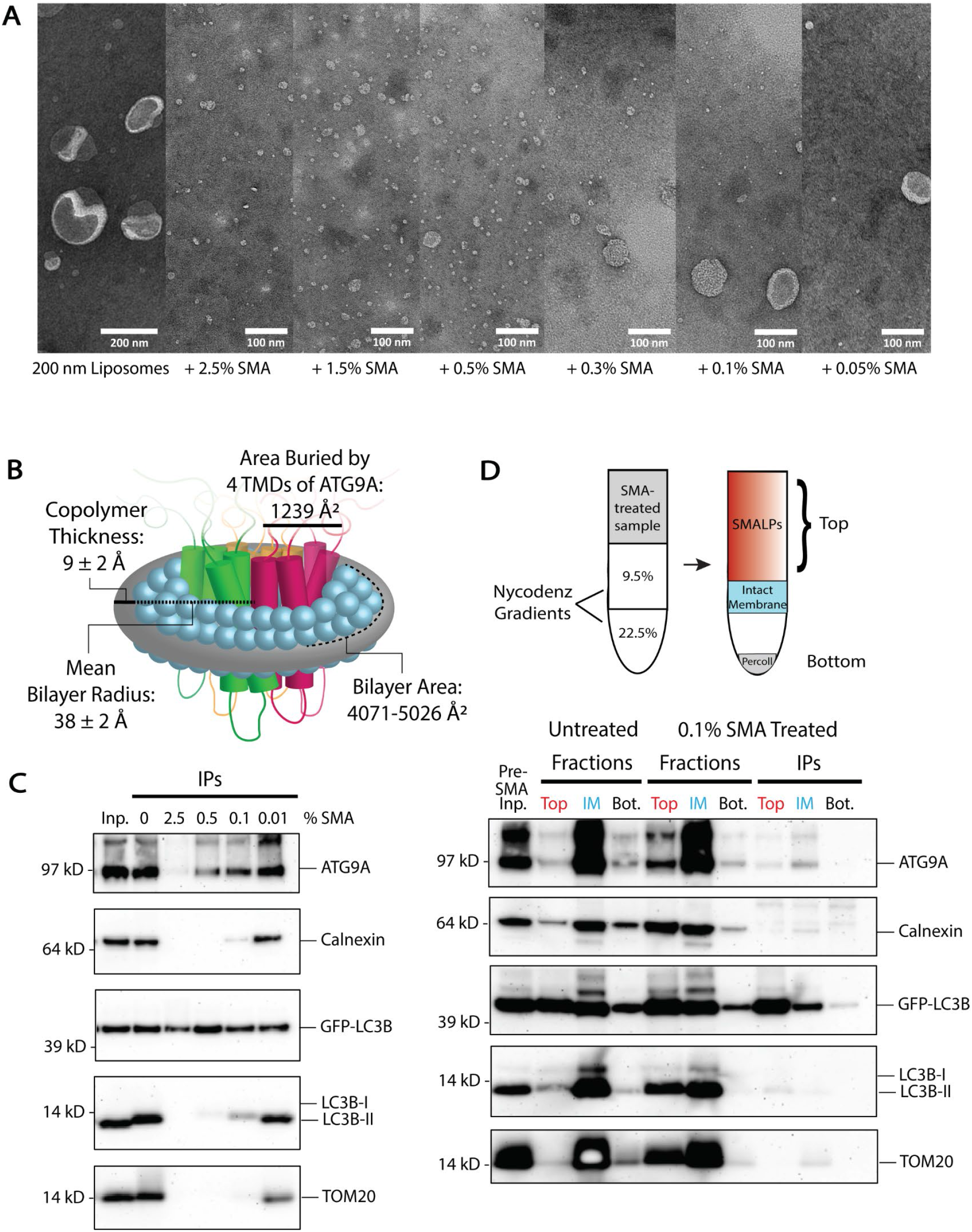
Styrene maleic acid (SMA) copolymer nanodiscs capture ATG9A and LC3B together in nanoscale membranes. **(A)** Transmission electron microscopy (TEM) of 200 nm liposomes treated with varying concentrations of SMA copolymer. The titration reveals the formation of SMALPs even at low SMA concentrations, with complete solubilization of liposomes at 1.5%-2.5% SMA. Scale bar in first panel: 200 nm. Scale bars in SMA-treated panels: 100 nm. **(B)** Cartoon of the transmembrane domains (TMDs) of the ATG9 trimer in a SMALP. The average 2:1 SMA copolymer nanodisc measures between 8.6 and 10.2 nm across, and can theoretically contain up to 40 transmembrane helices. SMALP measurements are from small angle neutron scattering (SANS) results (Jamshad et al., 2015), and the measured surface area buried by each 4-TMD bundle is from structural assessment (Guardia et al., 2020). **(C)** Immunoblot of a SMA titration on the pre-AP membrane fraction from ATG2 DKO HEK293 cells followed by immunoprecipitation (IP) of GFP-LC3B. ATG9A and GFP-LC3B co-IP with little to no membrane contaminants at higher concentrations of SMA. At lower concentrations of SMA, where membranes are incompletely solubilized into nanodiscs, the IP continues to pull-down all components. Input protein for each IP: 200 μg. Loaded input protein: 3 μg (1.5%). Loaded IPs: 5% of the total collected beads. **(D)** Immunoblot of the separation of intact membranes from SMALPs via density gradient fractionation (cartoon above) after treatment with a low concentration of SMA (0.1%) on the pre-AP membrane fraction from ATG2 DKO HEK293 cells. SMALPs float like cytosolic protein, while intact membranes sink to the 22.5% Nycodenz interface. The addition of 0.1% SMA drives each of the membrane associated proteins into the Top fraction relative to untreated controls, indicating the capture of these various proteins in SMALPs. After flotation, GFP-LC3B in the top fraction co-immunoprecipitates with ATG9A and LC3B-II, but not contaminants, confirming that GFP-LC3B SMALPs are largely limited to autophagy proteins. Input protein on gradient: 600 μg. Input for each IP: 90% of the material collected from the gradient. Loaded pre-SMA input protein: 3 μg (0.5%). Loaded gradient protein: 5% of the total collected. Loaded IPs: 10% of the total collected beads.

The capability of SMA to punch out sections of membrane and preserve the bilayer provides a mechanism to biochemically establish whether ATG9A and LC3B are present on the same membrane. ATG9A is likely a trimer in the membrane (Guardia et al., 2020; Maeda et al., 2020; Matoba et al., 2020). Each monomeric bundle of transmembrane domain α-helices buries 1,239 Å^2^ of surface area on the membrane (Guardia et al., 2020), thus an ATG9A trimer can be captured in an average-sized SMALP (~surface area between 4,071-5,026 Å^2^ (Jamshad et al., 2015)), with enough space left for lipids including those that may be conjugated to LC3B if both proteins are on the same span of membrane (Figure 4B).

To develop SMA as a tool to detect protein co-residency on natural membranes, SMA was titrated onto membranes enriched in the pre-AP fraction from ATG2 DKO cells, and then GFP-LC3B was immunoisolated. In the absence of SMA, GFP-LC3B immunoisolation pulls down essentially the same distribution of all proteins as the starting pre-AP fraction (Figure 4C, compare Inp. to 0), suggesting elements of this compartment remain organized and associated with the ER during the pull-down. SMA-treated immunoisolates contained both ATG9A and GFP-LC3B; at low SMA concentrations these immunoisolates harbor both SMALPs and incompletely solubilized membranes, and thus Calnexin and TOM20 are recovered. Crucially at higher concentrations of SMA, contaminating ER and mitochondrial membrane markers are lost, but GFP-LC3B and ATG9A continue to co-purify along with some detectable endogenous LC3B-II (Figure 4C) suggesting these molecules are co-resident in individual SMALPs.

High SMA concentrations fully solubilize membranes, potentially raising questions about the integrity of the membranes from which the individual SMALPs arise. At low SMA, each SMALP is more akin to a single bite from an apple and may better represent the natural protein distribution in the original membrane. To characterize SMALPs generated at low SMA concentrations, separate from the remaining largely intact membrane, we devised a density gradient membrane isolation that separates SMALPs from intact membranes. ATG2 DKO pre-AP fraction was treated with 0.1% SMA and applied to the same gradient as the first step in Figure 3A. We discovered that SMALPs efficiently solubilized transmembrane proteins into particles that collect at the top of the gradient while intact membranes appear to continue to concentrate at a lower interface (Figure 4D). Immunoisolation from this SMALP-containing top fraction showed co-purification of ATG9A, GFP-LC3B and some endogenous LC3B away from other membrane contaminants. Thus, immunoisolation with either high or low SMA percentages, capture nanomembrane expanses that are populated by both ATG9A and lipidated forms of LC3B.

### ATG9A and LC3B are co-resident on the autophagosome

We next used SMA isolations to compare how the co-residency of ATG9A and LC3B proteins varies between the pre-autophagosomal compartment in ATG2 DKO cells and the autophagosome-enriched fraction from WT cells (Figure 5, A-C). In both cases we found that immunoprecipitation of whole organelles (WO) without SMA polymer did not allow for significant purification of autophagy proteins away from contaminants. Immunoisolation of SMA-treated samples lead to the specific capture of ATG9A with GFP-LC3B. Interestingly, the ratio of ATG9A to GFP-LC3B was about 7 times lower in mature autophagosomes (Figure 5C; Figure S3). This suggests that as autophagosomes mature the surface density of ATG9A comes down relative to LC3B surface densities— precisely what would be expected if the membrane was expanding by lipid transfer which would facilitate more LC3B lipidation without introducing more ATG9A.

**Figure 5.**
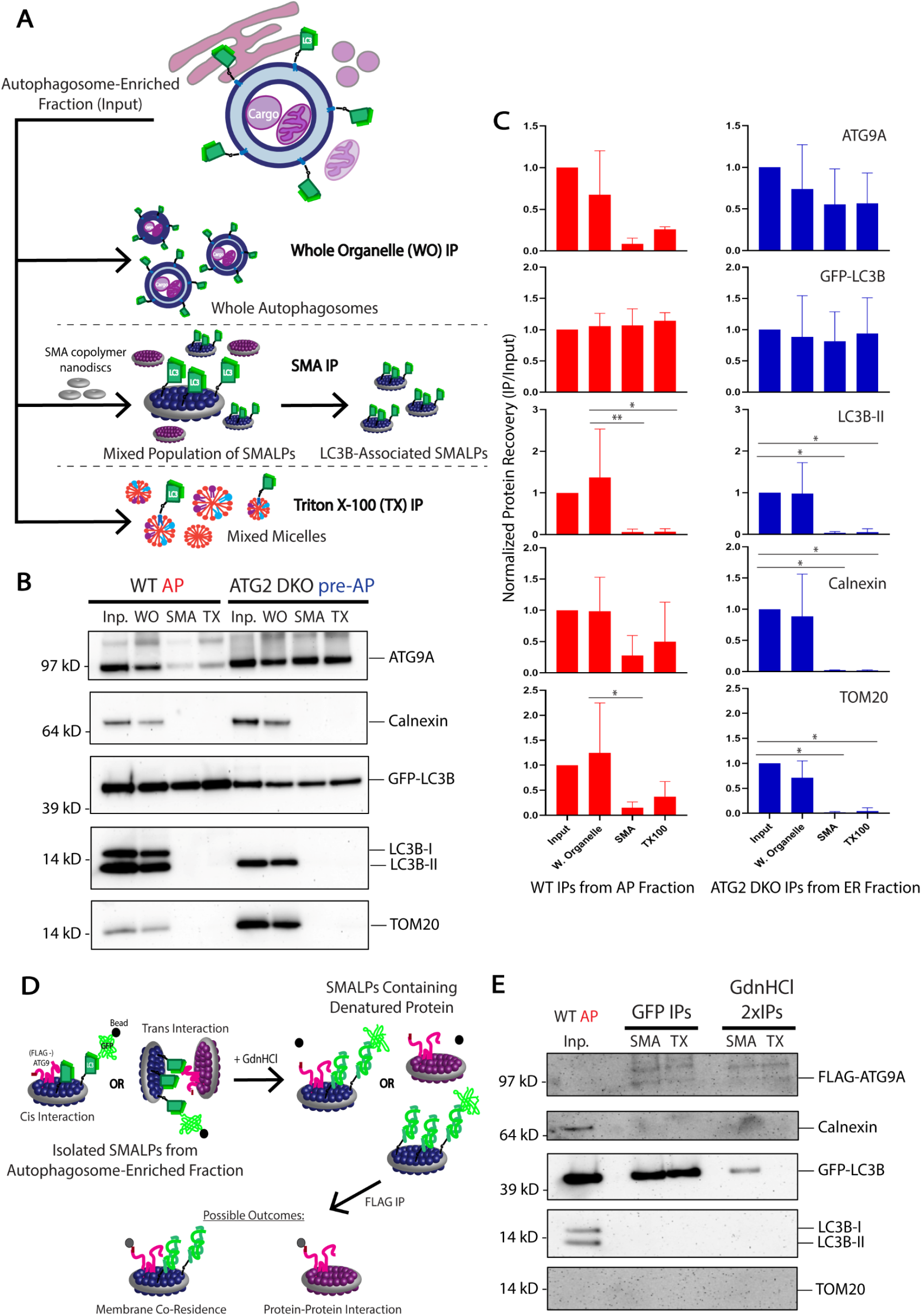
*Cis*-interacting ATG9A and LC3B are co-resident on autophagosomes and autophagosome precursor membranes. **(A)** Schematic cartoon outlining the three types of IPs performed on membrane fractions enriched for ATG9A and LC3B collected as in **Figure 3** from WT and ATG2 DKO HEK293 cells. These IPs were performed for both WT fraction 6 (autophagosome-enriched) and ATG2 DKO fraction 5 (pre-autophagosomal structure-enriched). **(B)** Immunoblot of IPs for GFP-LC3B from untreated fractions (whole organelle, WO), 12:1 SMA:protein (~2.5%) SMA-treated fractions (SMA), and 1% Triton X-100-treated fractions (TX). ATG9A and GFP-LC3B continue to co-IP in SMALPs as in **Figure 4**, but also after conventional detergent-based IP. Input protein for each IP: 50 μg. Loaded input protein: 3 μg (6%). Loaded IPs: 10% of the total collected beads. **(C)** Densitometric quantification of ATG9A, GFP-LC3B, LC3B-II, Calnexin, and TOM20 in each type of IP. The intensity of the bands in **B** were normalized to the input, and statistical significance was assessed by two-way ANOVA. *, adjusted p-value < 0.05. **, adjusted p-value < 0.01. Quantifications between cell lines are shown in **Figure S3**. **(D)** Schematic cartoon outlining a protein denaturation experiment between sequential IPs in order to separate SMALPs interacting *in trans*. ATG9A and GFP-LC3B are in the same SMALP from the same membrane only if they continue to co-purify after protein denaturation with guanidine hydrochloride (GdnHCl). **(E)** Immunoblot of sequential IPs, first with GFP-LC3B and then with FLAG-ATG9A, from double tagged WT HEK293 cells. After the first IP, the sample was treated with 6 M GdnHCl before diluting over 25X and performing a second IP against FLAG-ATG9A. Recovery of GFP-LC3B after the second IP is only detected in SMALPs, but not in soluble Triton-X micelles. Input protein for GFP IPs: 35 μg. Input for FLAG IPs: 98% of the material collected from the GFP IPs. Loaded input protein: 0.35 μg (1%). Loaded GFP IPs: 1% of the total collected from the beads. Loaded FLAG IPs: 50% of the total collected from the beads after GdnHCl treatment.

Because SMALPs carry only ~140 lipids, the capture of two separate proteins in a single disc is unlikely to occur at random. For example, on liposomes reconstituted with both GFP-LC3B-II and LC3B-II, the frequency of capturing both proteins is very low (Figure S3, J and K). ATG9A co-isolates with GFP-LC3 in SMALPs much more efficiently than would be expected if they were randomly distributed and furthermore, ATG9A and GFP-LC3B also co-isolate together in a classic Triton X-100 mediated detergent immunoisolation (Figure 5, B and C). This suggests these proteins either directly interact or both participate in a larger protein complex. A preprint article has eluded to such an interaction (Zhang et al., 2020), but an ATG9-LC3 binding interaction did not surface in a previous proteomic study (Behrends et al., 2010). Formally, such a complex could form between membranes rather than on a single membrane and thus it became important to determine whether the interaction was *cis* on one SMALP or *trans* on two SMALPs. We employed a protein denaturation protocol to disrupt any protein-protein interactions that could support a multi-SMALP *trans* complex (Figure 5D). We created WT and ATG2 DKO HEK293 cell lines double-labeled with GFP-LC3B and FLAG-ATG9A, and enriched for autophagosomes or pre-autophagosomal site membranes, respectively, as shown in Figure 3. Samples were then treated with either SMA or Triton X-100 and immunoisolated for GFP-LC3B before being treated with guanidine hydrochloride (GdnHCl) to denature possible protein interactions. Treatment with 6 M GdnHCl did not change the buffered neutral pH of the sample solution, importantly maintaining favorable conditions for the pH-sensitive SMA copolymer (Dörr et al., 2016; Scheidelaar et al., 2016). After diluting the samples containing GdnHCl at least 25 times the samples were put through a second round of immunoisolation targeting FLAG-ATG9A. As expected for a protein-protein complex, FLAG-ATG9A and GFP-LC3B were no longer co-precipitated in the detergent treated sample (Figure 5E). In contrast, FLAG-ATG9A continued to pull-down GFP-LC3B in the SMA treated samples (Figure 5E), as expected if they were each independently anchored in the same nanomembrane.

Collectively, this work demonstrates biochemically that mammalian ATG9A is incorporated onto the autophagosome membrane. Furthermore, in the absence of ATG2-mediated membrane expansion, LC3B still finds ATG9A vesicles at a site proximal to the ER and undergoes lipid conjugation. Taken together, this data suggests that ATG9 vesicles are the seed membrane for the autophagosome.

## Discussion

ATG9 vesicles are required for autophagosome formation, but precisely where they function has been challenging to establish (Feng and Klionsky, 2017; He et al., 2008; He et al., 2006; Mari et al., 2010; Orsi et al., 2012; Yamamoto et al., 2012; Young et al., 2006). In particular, in mammals the majority of ATG9 vesicles are not localized to the site of autophagosome biogenesis at any given time, and fluorescence imaging during autophagosome biogenesis appeared to favor a peripheral rather than integrated role for these vesicles as autophagosomes grew (Claude-Taupin et al., 2021; Mailler et al., 2021; Mattera et al., 2017; Orsi et al., 2012; Puri et al., 2013; Young et al., 2006). In contrast, in yeast, biochemistry proved to be a valuable approach to capture Atg9 within autophagosomes (Yamamoto et al., 2012). Here, we have re-engineered the structural biology tool styrene-maleic acid nanodiscs to develop a nanoscale tool for capturing proteins jointly embedded in a common membrane and used this device to formally establish that both ATG9A and LC3B-II reside in the same membranes, both in purified autophagosomes and in ER-rich membranes replete with pre-autophagosomal material. Further, we show that in the absence of the lipid transport protein ATG2, ATG9 vesicles collect near the ER but remain competent to collect lipidated LC3B. How the vesicle cluster is maintained in its organization is unclear. It has been reported that p62 is capable of phase separation both *in vitro* and *in vivo* (Sun et al., 2018), so it is possible that p62 provides a matrix that holds this compartment together. Interestingly, ATG9A vesicles can also be driven into similar large phase-separated clusters by proteins regulating synaptic vesicle organization (Park et al., 2022) suggesting they may generally be substrates for these types of adaptors. Given the composition of the pre-ATG2 compartment, we propose that it captures the initiation stage of autophagosome biogenesis before membrane expansion can occur and by extension traps the seed membrane for the autophagosome. These results are consistent with many other studies placing autophagosome biogenesis at contact sites involving the ER (Gómez-Sánchez et al., 2018; Hamasaki et al., 2013; Maeda et al., 2019; Osawa et al., 2019; Tang et al., 2019; Valverde et al., 2019).

Our findings offer one of the final pieces to the phagophore half of the lipid transport facilitated expansion model of autophagosome biogenesis (Figure 1A and (Ghanbarpour et al., 2021; Maeda et al., 2020; Matoba et al., 2020; Melia and Reinisch, 2022; Noda, 2021)). ATG2 binds the phagophore, where it is held in position by peripheral protein receptors including GABARAP/L1 and WIPI4 (Bozic et al., 2020; Chowdhury et al., 2018; Gómez-Sánchez et al., 2018; Kotani et al., 2018; Maeda et al., 2019). Our data presented here puts ATG9 also on the phagophore membrane, where its scramblase function can populate the inner leaflet of the growing organelle as it is fed lipid by ATG2. Various studies indicate that ATG2 proteins directly engage ATG9 (Ghanbarpour et al., 2021; Gómez-Sánchez et al., 2018) forming a scramblase-transporter complex. Intriguingly, the other side of ATG2 is bound to the ER through interactions that include additional scramblases (Ghanbarpour et al., 2021) suggesting there may be a broad benefit to coupling bulk lipid transport with scrambling activity. Future work must still define the driving force behind unidirectional lipid flow through ATG2 in order to grow the autophagosome.

## Experimental Procedures

### Plasmids and Reagents

1,2-Dioleoyl-*sn*-glycero-3-phosphoethanolamine (DOPE; 850725C), 1-palmitoyl-2-oleoyl-*sn*-glycero-3-phosphocholine (POPC; 850457C), l-α-phosphatidylinositol (from bovine liver – blPI; 840042C), 1,2-dioleoyl-*sn*-glycero-3-phosphocholine (DOPC; 850375C), 1,2-dioleoyl-*sn*-glycero-3-phosphoethanolamine-N-(lissamine rhodamine B sulfonyl)(Rhod-PE; 810150C) were purchased from Avanti Polar Lipids. ATP (AB00162), DTT (AB00490), isopropyl β-d-thiogalactopyranoside (AB00841), NaCl (AB01915), sucrose (AB01900), Tris HCl (AB02005) were purchased from AmericanBio. 100% glycerol, anhydrous (2136), CaCl_2_ (1332), EDTA (8993), MgCl_2_ (2444) were purchased from J. T. Baker. 0.05% Trypsin-EDTA 1X (25300054), Dulbecco’s phosphate buffered saline (DPBS) 10X (14200075), DMSO (D2650) were purchased from Thermo Fisher Scientific. Borosilicate glass tubes 10 X 75 mm (47729568) were purchased from VWR.

For transient rescue of the ATG9A KO cell line, 3xFLAG-ATG9A was subcloned from pCMV-10-3xFLAG-ATG9A (described in (Ghanbarpour et al., 2021)) into pLVX-puro (Clontech; 632159) for stable cell line creation. pLVX-GFP-LC3B was generated by Gibson assembly using pLVX-Puro backbone, and gel-purified PCR products of GFP-6xGS linker and LC3B to form the GFP-LC3B insert. GFP-LC3B was then cut from the pLVX-puro backbone using EcoRI and BamHI restriction sites, and inserted into the digested and gel-purified backbone of pLVX-EGFP-IRES-Neo from R. Sobol (University of Pittsburgh School of Medicine, Pittsburgh, PA; Addgene; 128660) for double-labeled stable cell line creation.

Antibodies used for immunofluorescence in this study include anti-LC3B (MBL; PM036; rabbit; 1:500), anti-LC3B, clone 2G6 (Nanotools; 0260100; mouse; 1:100), anti-ATG9A (Abcam; ab108338; rabbit; 1:250), anti-p62 (BD Biosciences; 610832; mouse; 1:250), anti-WIPI2, clone 2A2 (Millipore; MABC91; mouse; 1:250), anti-RB1CC1 (Thermo Fisher Scientific; 172501AP; rabbit; 1:100), Alexa Fluor 405 goat anti-rabbit IgG (Thermo Fisher Scientific; A31556), Alexa Fluor 405 goat anti-mouse IgG (Thermo Fisher Scientific; A31553), Alexa Fluor 488 goat anti-rabbit IgG (Thermo Fisher Scientific; A11008), Alexa Fluor 594 goat anti-rabbit IgG (Thermo Fisher Scientific; A11037), and Alexa Fluor 594 donkey anti-mouse IgG (Thermo Fisher Scientific; A21203). Antibodies used for immunoblotting in this study include anti-LC3B (Cell Signaling Technology; 3868S; rabbit), anti-GFP (Cell Signaling Technology; 2956S; rabbit), anti-ATG9A (Abcam; ab108338; rabbit), anti-Calnexin (BD Biosciences; 610524; mouse), anti-TOM20 (Cell Signaling Technology; 42406S; rabbit), anti-p62 (BD Biosciences; 610832; mouse), anti-GAPDH (Abcam; ab9484; mouse), ECL anti-rabbit IgG horseradish peroxidase-linked (GE Healthcare; NA934V), and ECL anti-mouse IgG horseradish peroxidase-linked (GE Healthcare; NA931V).

### Cell culture

HEK293 cells were cultured at 37°C and 5% CO_2_ in DMEM (Thermo Fisher Scientific; 11965092) supplemented with 10% FBS (Thermo Fisher Scientific; 10438062) and 1% penicillin-streptomycin (Thermo Fisher Scientific; 15140122). For experiments to collect autophagosome membranes, cells were starved by incubation in Earle’s Balanced Salt Solution (Thermo Fisher Scientific; 24010043) and treated with 0.1 μM Bafilomycin A1 (Enzo; BML-CM110-0100) for 4-6 hrs.

### Lentivirus production and transduction

HEK293 cells were seeded into a 10-cm plate. At 70% confluence, cells were transfected with 4.3 μg psPAX2 (Addgene; 12260), 0.43 μg pCMV-VSV-G (Addgene; 8454), and 4.3 μg target plasmid using 36 μL Lipofectamine 3000 (Thermo Fisher Scientific; L3000008). DNA was added to 500 μL Opti-MEM (Thermo Fisher Scientific; 31985070) and Lipofectamine 3000 to 500 μL Opti-MEM in separate 1.5 mL microcentrifuge tubes. The tubes were then mixed and left for 15 min before adding the mixture dropwise to cells in fresh DMEM. After overnight incubation at 37°C, the medium was replaced with fresh DMEM. Cell medium was then collected every 24 hrs for 2 days and filtered with a 0.45-μm syringe filter (Pall; 4184). Generated virus was stored at 4°C overnight the first day to pool both collections, and was then either used immediately or was aliquoted and stored at −80°C.

HEK293 cells to be transduced were seeded into a 6-well plate. Undiluted virus was added dropwise to cells with 10 μg/mL polybrene (Sigma-Aldrich; TR1003G), and incubated at 37°C for 24 hrs. The medium was replaced with fresh DMEM and left for another 24 hrs. The cells were then treated with 2 μg/mL puromycin (Clontech; 631306) under selection for at least 1 week. In the case of double-labeled cells, the process was completed again and selection was performed by treating cells with 2 mg/mL Geneticin (Thermo Fisher Scientific; 10131035) for at least 2 weeks.

### Focused Ion Beam Scanning Electron Microscopy and Correlative Light Electron Microscopy

GFP-LC3B over-expressed ATG2 DKO HEK293 cells were plated on 35 mm MatTek dish (P35G-1.5-14-CGRD). Cells were pre-fixed in 4% PFA + 0.25% glutaraldehyde then washed before fluorescence light microscopy imaging. Regions of interest were selected and their coordinates on the dish were identified using phase contrast. Cells were further fixed with 2.5% glutaraldehyde in 0.1 M sodium cacodylate buffer, postfixed in 2% OsO4 and 1.5% K4Fe(CN)6 (Sigma-Aldrich) in 0.1 M sodium cacodylate buffer, en bloc stained with 2% aqueous uranyl acetate, dehydrated, embedded in Embed 812, polymerized at 60 °C for 48 hrs.

Epon block was glued onto the SEM sample mounting aluminum stub and platinum en bloc coating on the sample surface was carried out with the sputter coater (Ted Pella, Inc., Redding, California). The cell of interest was relocated under SEM imaging based on pre-recorded coordinates and FIB-SEM imaged in a Crossbeam 550 FIBSEM workstation operating under SmartSEM (Carl Zeiss Microscopy GmbH, Oberkochen, Germany) and Atlas 5 engine (Fibics Inc., Ottawa, Canada). The imaging resolution was set at 7 nm/pixel in the X, Y axis with milling being performed at 7 nm/step along the Z axis to achieve an isotropic resolution of 7nm/voxel. Images were aligned and exported with Atlas 5 (Fibics Inc., Ottawa, Canada), further processed and analyzed with DragonFly Pro software [Object Research Systems (ORS) Inc., Montreal, Canada]. 3D segmentation was also performed using DragonFly Pro software.

### High-pressure freezing, freeze substitution, and Transmission Electron Microscopy (TEM)

ATG2 DKO HEK293 cells were cultured on 3 mm sapphire discs (TechnoTrade International, Inc.) and flash frozen with a Leica EM HPM100 machine in 20% BSA (Sigma-Aldrich). Frozen cells were transferred to liquid nitrogen for storage. For freeze substitution, the cells on the sapphire discs were transferred to a Leica EM AFS2 apparatus, where they were first immersed in acetone containing 1% OsO4, 1% glutaraldehyde, 0.1% uranyl acetate, and 1 % H2O at −90 °C for 3-6 hrs. Subsequently, they were shifted to 0 °C within 12 hrs. Samples were then fixed in fresh 1% OsO4 and 1% H2O in acetone for 1 hr at room temperature (RT), washed in pure acetone (10 min x 3) and infiltrated with epon using the following sequence: 1 hr epon/acetone 1:1 (v/v); 1 hr pure epon by 2 times at RT. Finally, embedded samples were polymerized at 60 °C for 48 hrs. Ultrathin sections (60 nm) were observed in a Philips CM10 microscope at 80 kV and images were taken with a Morada 1kx1k CCD camera (Olympus). Except noted all reagents were from EMS (Electron Microscopy Sciences, Hatfield, PA).

### Immunofluorescence and confocal microscopy

HEK293 cells were seeded onto coverslips in a 24-well plate. If cells were treated with Bafilomycin A1, 0.1 μM was used for 2 hrs. Cells were rinsed in PBS, fixed in 4% PFA (Electron Microscopy Sciences; 15710), washed three times with PBS, and permeabilized in ice-cold methanol by dipping the coverslip in it 20 times followed by blocking in 3% BSA (Sigma-Aldrich; A9647) in PBS for 15 min (**Figures 2 A-H and S1**). Alternatively, cells were permeabilized and blocked in 0.1% saponin (Sigma-Aldrich; 47036) and 3% BSA in PBS instead of methanol permeabilization (**Figures 2I and S2 A**). Cells were then incubated in primary antibody at indicated concentrations (see Reagents) overnight at 4°C. Methanol-permeabilized cells were washed three times in PBS and saponin-permeabilized cells were washed three times in PBS containing 0.1% saponin and 3% BSA. Secondary antibody was applied in a 1:600 dilution for 1 hr at room temperature in the dark. Cells were washed again three times, and mounted on pre-cleaned microscope slides with ProLong Gold antifade reagent (Thermo Fisher Scientific; P36934).

Imaging was performed at the Center for Cellular and Molecular Imaging Facility at Yale. The 63x oil-immersion objective was used on an inverted Zeiss LSM 880 laser scanning confocal microscope with AiryScan, using Zen acquisition software.

### Generation of ATG9A knockout using CRISPR/Cas9

CRISPR guide RNAs (gRNAs) targeting the third, sixth, seventh, and eighth exons of ATG9A were accessed online from the Toronto KnockOut Library (University of Toronto; TKOv3, (Hart et al., 2017)). The final target sequences were 5’-TATAGGAGGCCTCTAGGCGC-3’ (TKO1), 5’-TAGTGAAGGCAACCACAAAG-3’ (TKO2), 5’-GAAGCTGTCTTCTTCACCCG-3’ (TKO3), and 5’-AGATGAACTTGATAAGCCGG-3’ (TKO4). Complementary sequence to the overhangs created in the vector backbone after digestion were added to the 5’ ends of the forward and reverse gRNA primers. The gRNAs were cloned into pX458 (Addgene; 48138) using BpiI (Thermo Fisher Scientific; FD1014) as described previously (Ran et al., 2013). HEK293 cells were transiently transfected with the constructs containing gRNAs using Lipofectamine 3000 according to the manufacturer’s suggestion. After 48 hrs, cells were sorted into a 96-well plate for single clones using a BD FACSAria cell sorter at the Flow Cytometry Facility at Yale. The cells were validated by genotyping, Western blotting, and immunofluorescence. For genotyping, briefly, genomic DNA from single clones was extracted using QuickExtract DNA extraction solution (Lucigen; QE0905T) according to the manufacturer’s suggestion. PCR products were generated using the following surveyor primers: TKO1-F, 5’-CAGGAAAGCAGCAGTAGACAC-3’; TKO1-R, 5’-GAGACATTAAAGGTCCCAGAG-3’; TKO2-F, 5’-GTAGTGGTGGCTGAGGTTACC-3’; TKO2-R, 5’-CTTGGCAGAGACGCTGCTATC-3’; TKO3-F, 5’-GAGGCAACAACCCCACCTTC-3’; TKO3-R, 5’-CCATAGAGTGACCAGCAGCG-3’; TKO4-F, 5’-CTTGAGTTCTGCTAGCAGGTG-3’; TKO4-R, 5’-GACCCCTTTGCCCTATATTAG-3’. Surveyor primers were designed to be unique to the target gene using NCBI Primer-BLAST and tested on WT HEK293 cell genomic DNA for easily excisable bands from a 1% agarose gel before using them on ATG9A KO candidate genomic DNA. The 400-600 bp PCR products were gel purified and sequenced. The chosen ATG9A KO HEK293 cell line was cut with gRNA TKO1, creating an in-frame stop codon after 36 bases making the expressed ATG9A peptide fragment only 12 amino acids long.

### Density gradient membrane fractionation

Membrane fractionation was modified from (Strømhaug et al., 1998). In summary, WT HEK293 cells were seeded into 60 15-cm plates, ATG2 DKO HEK293 cells (Valverde et al., 2019) were seeded into 40 15-cm plates, and both cell lines were grown to 80-90% confluency. WT cells were starved in EBSS and treated with 0.1 μM Bafilomycin A1 for 4-6 hours before harvesting to accumulate autophagosomes, and ATG2 DKO cells were collected untreated (autophagy factors accumulate without intervention, (Valverde et al., 2019)). Cells were scraped from the plates, resuspended in 1 mL sucrose homogenization buffer (SHB; 50 mM Tris pH 7.4, 150 mM NaCl, 10% sucrose, and 1X EDTA-free protease inhibitor cocktail [Sigma-Aldrich; 11873580001]), and lysed with a 2-mL Dounce homogenizer (Sigma-Aldrich; DWK8853000001). The lysates were centrifuged at 4000 rpm for 2 min at 4°C to pellet the nuclei and collect the post-nuclear supernatant (PNS). PNS fractions were loaded onto a Nycodenz (Accurate Chemical and Scientific Corp; 1002424) gradient (110 μL of 22.5% Nycodenz on bottom, 270 μL of 9.5% Nycodenz layered next, and 225 μL of PNS on top) in a thin-walled ultracentrifuge tube (Beckman Coulter; 355090) and centrifuged at 38,600 rpm with an SW55 rotor (Beckman Coulter; 342194) for 1 hr at 4°C. Fractions collected are shown in **Figure 3A**, and were recovered in the following volumes: F1, 50 μL; F2, 50 μL; F3, 80 μL; F4, 30 μL; F5/6, 120 μL. Fraction 5/6 (found at the interface between 22.5% and 9.5% Nycodenz and enriched in ER and autophagosome membranes (Strømhaug et al., 1998)) was diluted 1.25X and loaded on a Percoll (Sigma-Aldrich; P4937)/Nycodenz gradient (110 μL of 22.5% Nycodenz on bottom, 300 μL of 33% Percoll layered next, and 200 μL of Fraction 5/6 on top) in an ultracentrifuge tube and centrifuged at 27,600 rpm for 30 min at 4°C. The two fractions were recovered in the following volumes: F5, 80 μL; F6/Percoll, 80 μL. For every 240 μL of F6/Percoll collected, 168 μL of 60% (w/v) OptiPrep density gradient medium (Sigma-Aldrich; D1556) was added to the fraction. This membrane/Percoll/OptiPrep sample was loaded on the final OptiPrep gradient (408 μL of F6/Percoll/OptiPrep on bottom, 75 μL 30% OptiPrep layered next, and 120 μL of 1X SHB on top) in an ultracentrifuge tube and centrifuged at 27,300 rpm for 30 min at 4°C. Fraction 6 free of Percoll was collected in a volume of 40 μL from the 30% OptiPrep/buffer interface. All gradient mediums were diluted with 5X SHB to final concentrations in 1X SHB in 10 mL. All fractions were collected and pooled from multiple gradients.

### Lysis, gel electrophoresis, immunoblotting, and Coomassie staining

Nearly all experiments required maintenance of membrane integrity, so cells were Dounce homogenized for lysis without detergent. For ATG9A KO screening and rescue, cells were lysed in 400 μL lysis buffer (50 mM Tris pH 7.4, 150 mM NaCl, 1% Triton X-100 [AmericanBio; AB02025], 1X EDTA-free protease inhibitor cocktail), incubated on ice for 5 min, centrifuged at 14,000 rpm for 10 min at 4°C, and the supernatant was collected and used immediately or stored at −20°C short term. Protein concentration was determined using protein assay dye reagent (Bio-Rad; 5000006) and a Bio-Rad SmartSpec Plus spectrophotometer. Samples were prepared in 1X LDS loading buffer (Thermo Fisher Scientific; NP0007) with 15 mM DTT, and then boiled at 98°C for 5 min. Samples were electrophoresed on 12% Bis-Tris precast gels (Thermo Fisher Scientific; NP0341BOX or NP0342BOX) with a SeeBlue Plus2 pre-stained protein standard (Thermo Fisher Scientific; LC5925), typically at 150 V for 75 min in 1X MOPS running buffer (Thermo Fisher Scientific; NP0001). For separation of GFP-LC3B-I from GFP-LC3B-II, samples were electrophoresed on 8% Bolt Bis-Tris Plus precast gels (Thermo Fisher Scientific; NW00080BOX) in 1X Bolt MOPS running buffer (Thermo Fisher Scientific; B0001), typically at 180 V for ~60 min to run the GFP band to the bottom of the gel. Gels were subsequently transferred to Immobilon-FL PVDF membranes (Sigma-Aldrich; IPFL00010) in 1X transfer buffer (Thermo Fisher Scientific; NP00061) containing 20% methanol (Sigma-Aldrich; 179337) at 30 V for 75 min.

For immunoblotting, membranes were blocked with 5% Omniblok (AmericanBio; AB10109) in PBST (PBS containing 0.1% Tween 20 [AmericanBio; AB02038]) for 1 hr at room temperature. Membranes were then rinsed with PBST, and incubated with primary antibody (see Reagents) diluted 1:1000 in PBST containing 5% BSA and 0.02% sodium azide (Sigma-Aldrich; S2002) overnight at 4°C. Membranes were washed three times in PBST before incubation with secondary antibody (Sigma-Aldrich; GENA931 or GENA934) diluted 1:5000 in PBST containing 5% Omniblok for 1 hr at room temperature. Membranes were then washed three times in PBST and treated with SuperSignal West Femto substrate (Thermo Fisher Scientific; 34096) for 5 min before imaging with the Bio-Rad VersaDoc imaging system.

For *in vitro* lipidation followed by IP, gels were electrophoresed in 1X MES running buffer (Thermo Fisher Scientific; NP0002), typically at 200 V for 55 min. The gels were stained for protein by incubation in Imperial Protein Stain (Thermo Fisher Scientific; 24615) for 1 hr at room temperature. Gels were then destained in water overnight at room temperature before imaging with the Bio-Rad VersaDoc imaging system.

### Densitometry and quantification

Densitometry quantifications of both immunoblots and Coomassie protein stained gels were performed using ImageJ software. For density gradient membrane fractionation (**Figures 3 and S3**), the band intensity of each fraction was normalized to that of the lysate, setting the lysate as 1. Mean values for WT HEK293 cell membrane fractions were compared to ATG2 DKO HEK293 cell membrane fractions, and statistical significance was determined by multiple unpaired t-tests. For immunoprecipitation (IP) experiments (**Figures 5 and S3**), the band intensity of each IP was normalized to that of the input, setting the input as 1. The mean values were first compared within cell type between different IP treatments, and statistical significance was determined by two-way ANOVA. The mean values were then compared between WT and ATG2 DKO values for each IP treatment, and statistical significance was determined by multiple unpaired t-tests. For *in vitro* lipidation followed by IP, the band intensity of LC3B-II was plotted as a ratio against GFP-LC3B to compare their recovery in each treatment. Statistical significance was determined by one-way ANOVA. Most data are n = 3, except for WT membrane fractions (**Figures 3 and S3**) that are n = 4 and WT Triton X-100 IPs (**Figures 5 and S3**) that are n = 2. All data were plotted with mean ± SD, and all significance values were considering adjusted p-value based on grouped analyses in Prism 9 (GraphPad). Asterisks indicate significance: *, p < 0.05; **, p < 0.01.

### Dialysis of styrene maleic acid

Styrene maleic acid (SMA) in NaOH (Polyscope; XIRAN SL30010P20) was dialyzed in SMA buffer (50 mM Tris pH 7.4 and 150 mM NaCl) before use in experiments. Briefly, 2 mL of SMA was injected into a 20 kD dialysis cassette (Thermo Fisher Scientific; 66003) and placed in 4 L of SMA buffer with gentle mixing for 5 hrs at room temperature. Buffer was replaced with 4 L of fresh SMA buffer and incubated with gentle mixing overnight at room temperature. The buffer was again changed, and the cassette incubated in 4 L of SMA buffer with gentle mixing for 5 hrs at room temperature. Dialyzed SMA was carefully removed from the cassette, and the percent concentration (w/v) was determined using an Abbe 3L refractometer (Bausch & Lomb). The calculation using refractive indices is as follows: (Sample_RI_ – Buffer_RI_) / 2 X 1000 = % SMA.

For *in vitro* lipidation followed by IP, ready-to-use SMA (Orbiscope; SMALP 30010P) was used instead of dialyzed stock.

### Liposome preparation

To prepare liposomes, lipids in chloroform were mixed in the noted compositions and dried to a thin film under nitrogen gas for ~10 min. The composition of liposomes for testing SMA (**Figure 4A**) was 30 mol percent DOPE, 60 mol percent POPC, and 10 mol percent PI. The composition of liposomes for *in vitro* lipidation assays (**Figure S3 J**) was 55 mol percent DOPE, 34.75 mol percent DOPC, 10 mol percent PI, and 0.25% rhodamine DOPE. The lipid film was further dried under vacuum for 1 hr. The lipids were reconstituted in SMA buffer (**Figure 4A**) or SN buffer (**Figure S3**; 20 mM Tris pH 8, 100 mM NaCl, and 5 mM MgCl2) and subjected to 7 cycles of flash freezing in liquid nitrogen and thawing in a 37°C bath. After freeze-thaw, liposomes were extruded with the LipSoFast-Basic extruder (Avestin) 21 times through 2 polycarbonate membranes (Sigma-Aldrich; WHA10417004 or WHA110405). For testing SMA, liposomes were extruded to 200 nm. For lipidation reactions, liposomes were first extruded to 100 nm and then were sonicated using the VirSonic 600 (VirTis) microtip sonicator to a size of 25 nm immediately prior to the lipidation reaction.

### Negative stain Transmission Electron Microscopy

For negative staining SMA-treated liposomes, 400 mesh Formvar carbon-coated copper grids (Electron Microscopy Sciences; FCF400CU) were glow discharged using a PELCO easiGlow glow discharge cleaning system (Ted Pella, Inc.). Samples were diluted 1:10 in buffer before being applied to the grid and stained with 2% uranyl formate. Imaging was performed on a JEOL JEM-1400 Plus microscope operated at 80 kV with a bottom-mount 4k×3k CCD camera (Advanced Microscopy Technologies).

### Styrene maleic acid membrane extraction, titration, and immunoprecipitation

SMA membrane extraction occurs spontaneously after addition to membranes (artificial or natural from cells). After addition of SMA in any concentration, samples were mixed and incubated for 5 min on ice to ensure complete extraction.

Titration was performed on 200 nm liposomes and on ATG2 DKO fraction 5 (**Figure 3**) due to plentiful membrane material. For liposome-based SMA titration, 50 mM liposomes were treated with 2.5%, 1.5%, 0.5%, 0.3%, 0.1%, and 0.01% SMA. They were then negatively stained as described previously. For SMA titration onto natural membranes, 200 μg of protein was treated with 2.5%, 0.5%, 0.1%, and 0.01% SMA, and incubated on ice for 5 min. The SMA to lipid ratios were about 12:1, 1:1.5, 1:3, and 1:33, estimating that the protein to lipid ratio in the ER is around 1:1. Autophagosomes have a lower protein to lipid ratio than the ER, so herein it was referred to as the SMA to protein ratio. Samples were then immunoisolated with GFP-LC3B using 50 μL of GFP microbeads from a GFP isolation kit (Miltenyi Biotec; 130091125) added to each sample. Samples were incubated with beads on ice for 2 hrs with gentle mixing every 15 min. The beads were then run through MACS 25 MS separation columns (Miltenyi Biotec; 130042201) on the MACS MultiStand magnetic stand (Miltenyi Biotec). Columns were primed with 500 μL 1X SHB, followed by the entire volume of samples containing beads, and then the beads were washed three times with 500 μL SHB. The beads were then eluted from the column by removing the column from the stand, holding it over a collection tube on ice, and adding 500 μL 1X SHB 100 μL at a time and allowing elution to occur by gravity. After the last 100 μL, any liquid left in the column was pushed through with a plastic plunger that comes with the column. Due to the large volume eluted, protein was precipitated from the beads using either methanol/chloroform precipitation (**Figure 4C**) or acetone precipitation (all other IP experiments).

For separation of whole organelles from SMA lipid particles (SMALPs) following treatment with low concentration SMA (**Figure 4D**), 600 μg of ATG2 DKO fraction 5 protein was treated with 0.1% SMA and incubated on ice for 5 min. SMA-treated sample and untreated control were loaded onto a Nycodenz gradient (110 μL of 22.5% Nycodenz on bottom, 270 μL of 9.5% Nycodenz layered next, and 100 μL of sample on top) in thin-walled ultracentrifuge tubes and centrifuged at 38,600 rpm for 1 hr at 4°C. The entire top fraction was carefully collected away from the visible intact membrane band (~400 μL), intact membranes were collected in 50 μL, as well as 50 μL from the bottom of the tube. The SMA-treated samples were then immunoisolated with GFP-LC3B using GFP microbeads as described previously.

SMA was used at a concentration of 2.5% per the literature. The SMA to protein ratio in using 2.5% SMA was calculated after the first set of IPs to be 12:1 SMA:protein, and was kept consistent for all following experiments.

### Three strategies for immunoprecipitation

For IPs against GFP-LC3B (**Figure 5B**), 50 μg of protein from either WT fraction 6 or ATG2 DKO fraction 5 (**Figure 3**) was either left untreated for whole organelle isolation, treated with a SMA to protein ratio of 12:1 SMA:protein (2.5%), or treated with 1% Triton X-100 for standard detergent-based IP (**Figure 5A**). Samples were incubated on ice for 5 min. All samples were then immunoisolated with GFP-LC3B using GFP microbeads as described previously.

### Methanol/chloroform protein precipitation

For precipitating protein from immunoisolated beads, 200 μL of methanol (Millipore; 106035) was added to 50 μL of sample (10% of the beads) and vortexed well. 50 μL of chloroform (Sigma-Aldrich; 650498) was then added, the sample was vortexed, 150 μL of ddH2O (Millipore; 115333) was added, the sample was vortexed again, and was then centrifuged at 14,000 x g for 2 min at room temperature. The top aqueous layer was removed, 200 μL of methanol was added to the sample, the sample was vortexed, and was then centrifuged at 14,000 x g for 3 min at room temperature. Methanol was removed carefully, and the sample was dried under vacuum for 2 min. The pellet was then resuspended in 1X LDS, boiled at 98°C for 10 min, centrifuged at 14,000 rpm for 10 min, beads were removed on a magnetic stand, and finally 15 mM DTT was added. Samples were then used for SDS-PAGE and immunoblotting.

### Acetone protein precipitation

For precipitating protein from immunoisolated beads, four times the sample volume of ice-cold acetone was added to 100 μL of sample (20% of the beads) and vortexed well. Acetone samples were incubated at −20°C for 60 min. Samples were then centrifuged at 14,000 rpm for 10 min at 4°C. Acetone was removed carefully, and samples were either dried uncapped under the fume hood for 30 min or under vacuum for 2-5 min. The pellet was then resuspended in 1X LDS, boiled at 98°C for 10 min, centrifuged at 14,000 rpm for 10 min, beads were removed on a magnetic stand, and finally 15 mM DTT was added. Samples were then used for SDS-PAGE and immunoblotting.

### Protein denaturation using guanidine hydrochloride and sequential immunoprecipitation

To make the determination between *cis* or *trans* protein-protein interaction (**Figure 5E**), WT FLAG-ATG9A expressing HEK293 cells were transduced with GFP-LC3B to create a double-labeled (DL) cell line for sequential IP. 35 μg of DL WT fraction 6 protein was treated with either SMA in a ratio of 12:1 SMA:protein or 1% Triton X-100 and immunoisolated with GFP-LC3B using 100 μL GFP microbeads, otherwise as described previously before protein precipitation. The eluted beads were then treated with 6 M guanidine hydrochloride (GdnHCl; [Thermo Fisher Scientific; 24115]) and incubated for 2 hrs at room temperature with mixing. Beads were removed by running the sample through a second MACS 25 MS separation column and collecting the eluate. The sample was then diluted to 225 mM GdnHCl in 1X SHB and applied to pre-washed Pierce anti-FLAG magnetic agarose beads (Thermo Fisher Scientific; A36797) in 50 mL tubes for 30 min at room temperature with mixing. The 50 mL tubes were centrifuged at 400 rpm to pellet the beads, and the flow-through was removed. The beads were then transferred to 1.5 mL microcentrifuge tubes and washed three times with 1X SHB on a magnetic stand. The protein was eluted by adding 1X LDS, boiling the samples at 98°C for 10 min, removing the beads on a magnetic stand, and adding 15 mM DTT. Samples were then used for SDS-PAGE and immunoblotting.

It is important to note that the dominant mechanism by which GdnHCl denatures protein is reported to be electrostatic interactions between the guanidinium ion and charged residues as well as the peptide backbone (O’Brien et al., 2007). For this reason, GdnHCl should unfold proteins without threatening the captured lipids or SMA copolymer itself. The SMA copolymer is reportedly pH-sensitive (Dörr et al., 2016), and the aqueous GdnHCl solution is weakly acidic. SMALP samples were buffered at pH 7.4 and the addition of GdnHCl to a final concentration of 6 M only shifted the pH to 7.2, remaining in the neutral range. In any case, SMA 2:1, which is the copolymer ratio we use for this study, functions at the widest pH range and remains soluble at a pH as low as 5.0 (Scheidelaar et al., 2016).

### Recombinant protein expression and purification

Mouse ATG3, human LC3B, and human GFP-LC3B were expressed and purified as previously described (Kauffman et al., 2018; Nath et al., 2014). For all *in vitro* experiments, LC3B and GFP-LC3B refer to protein expressed and purified from constructs ending at the reactive glycine (G120) of LC3B, and thus no ATG4-mediated pre-processing was needed for lipidation. In summary, human LC3B and GFP-LC3B were cloned in the pGEX-2T GST vector, and mouse ATG3 was cloned into pGEX-6p GST vector. All three proteins were expressed in BL21-Gold (DE3) competent cells (Agilent Technologies; 230132). The cells were cultured in 4 L of Luria Bertani broth (LB) medium with 50 mg/mL carbenicillin (Sigma-Aldrich; C1389) and induced with 0.5 mM isopropyl β-d-thiogalactopyranoside (IPTG). Bacterial pellets were treated with EDTA-free protease inhibitor tablets in either thrombin buffer (20 mM Tris pH 7.5, 100 mM NaCl, 5 mM MgCl_2_, 2 mM CaCl_2_, 1 mM DTT) for LC3B and GFP-LC3B or preScission protease buffer (50 mM Tris pH 7.5, 150 mM NaCl, 1 mM EDTA, 1 mM DTT) for ATG3. The cells were broken in a cell disrupter, and the lysate was incubated with pre-washed glutathione agarose beads (Sigma-Aldrich; G4501) for 3 h at 4°C. The beads were washed three times and then incubated with thrombin (for LC3B/GFP-LC3; 10 μL of thrombin [Sigma-Aldrich; T6884], 500 μL of thrombin buffer, 0.5 μL of DTT, 500 μL of beads) or preScission protease (for ATG3; 25 μL of preScission protease [Sigma-Aldrich; GE27084301], 500 μL of preScission protease buffer, 0.5 μL of DTT, 500 μL of beads) to cut the proteins from GST tags overnight at 4°C. Purified proteins were stored in 20% glycerol at −80°C.

### Protein expression and purification for human ATG7

Human ATG7 was expressed and purified as previously described (Choy et al., 2012). In summary, the ATG7-containing plasmid was transformed into Bacmid DNA. 2 μg of DNA was used to infect 8 X 10^5^ SF9 cells by using Cellfectin II (Thermo Fisher Scientific; 10362100) two times to increase viral titer to 5 X 10^6^ plaque-forming units/mL. 1 X 10^8^ plaque-forming unit/mL of SF9 cells were infected with virus and grew for 72 hrs. The cells were treated with PIC tablets in the lysis buffer (20 mM Tris pH 8, 500 mM NaCl, 20 mM imidazole, 1 mM DTT, 10% glycerol), sonicated with the Virsonic 600 (VirTis) microtip sonicator for 3 min (30 sec on, 30 sec off; intensity 3.5), and centrifuged at 18,000 rpm for 1 hr. The lysate was incubated with 1 mL of nickel resin (nickel–nitrilotriacetic acid–agarose) for 2 hrs at 4°C. The beads were washed with the wash buffer (20mM Tris pH 8, 300 mM NaCl, 20mM imidazole [Sigma-Aldrich; I5513], 1 mM DTT) three times and eluted with the elution buffer (20 mM Tris pH 7.5, 300 mM NaCl, 500 mM imidazole, 1 mM DTT). Purified proteins were stored in 20% glycerol at −80°C.

### Lipidation assay, float up, and immunoprecipitation following SMA extraction

Recombinant LC3B and GFP-LC3B underwent the lipidation process to be coupled to liposomes at the same time. In short, LC3B protein (30 μM), GFP-LC3B protein (15 μM), ATG3 (16 μM), ATG7 (4 μM), and sonicated liposomes (8 mM) were mixed with DTT (1 mM) in SN buffer. Lipidation was initiated by adding ATP (1 mM), the reaction was incubated at 37°C for 1 hr, 1 mM of ATP was added again to the reaction, and the reaction was incubated at 37°C for 1 hr.

Protein-coupled liposomes were floated up by loading them onto Nycodenz gradients (150 μL of lipidation reaction mixed with 150 μL of 80% Nycodenz [40% Nycodenz layer] on bottom, 250 μL of 30% Nycodenz layered next, and 50 μL of 1X SN buffer on top) in a thin-walled ultracentrifuge tube (Beckman Coulter; 355090), and centrifuging them at 48,000 rpm for 4 hrs at 4°C. Protein-coupled liposomes were recovered in 40 μL from the interface between 30% Nycodenz and buffer. Nycodenz was diluted with 5X SN buffer to final concentrations in 1X SN buffer in 10 mL. Protein-coupled liposomes were collected and pooled.

To validate natural membrane results with these liposomes decorated with only LC3B and GFP-LC3B, 37.5 μL of lipidated material was treated with 2.5% SMA, 0.5% SMA, or 1% Triton X-100. Samples were immunoprecipitated with GFP-LC3B using GFP microbeads as described previously.

## Acknowledgements

This work was supported by grants from the National Institutes of Health (R01 GM100930 to T.J. Melia; R01 GM141192 to K. Gupta; R01 NS036251 and P30 DK045735 to P. De Camilli), predoctoral training grants from the China Scholarship Council to S. Yu and from the National Institutes of Health to T.J. Olivas (T32 GM007223), and a Ruth L. Kirschstein National Research Service Award Individual Predoctoral Fellowship to Promote Diversity in Health-Related Research from the National Institutes of Health to T.J. Olivas (F31 GM142294). We are grateful to the facilities that supported this work including the Yale Center for Cellular and Molecular Imaging (both the fluorescence and electron microscopy facilities) and the Yale Diabetes Research Center. We also thank Chenxiang Lin for access to the transmission electron microscope at the Yale West Campus.

The authors declare no competing financial interests.

## Author contributions

T.J. Olivas designed and carried out the immunofluorescence experiments including creation of the ATG9A KO, density gradient membrane fractionations, and all assays using SMA copolymer nanodiscs except for performance of the SMA isolations from liposomes carried out by P. Choi. S. Nag prepared the reconstituted proteins for the lipidation of the liposomes in that experiment. S. Yu discovered the pre-ATG2 compartment in ATG2 DKO cells and initiated the collaboration with Y. Wu who performed the FIB-SEM, TEM and following electron microscopy analysis. L. Luan performed the initial 3D segmentation of the FIB-SEM results, later updated by Y. Wu. T.J Melia designed the project and supervised it with collaboration from P. De Camilli and K. Gupta. The manuscript was prepared by T.J. Melia and T.J. Olivas with input from all authors.

---

**Supplemental Figure 1.**
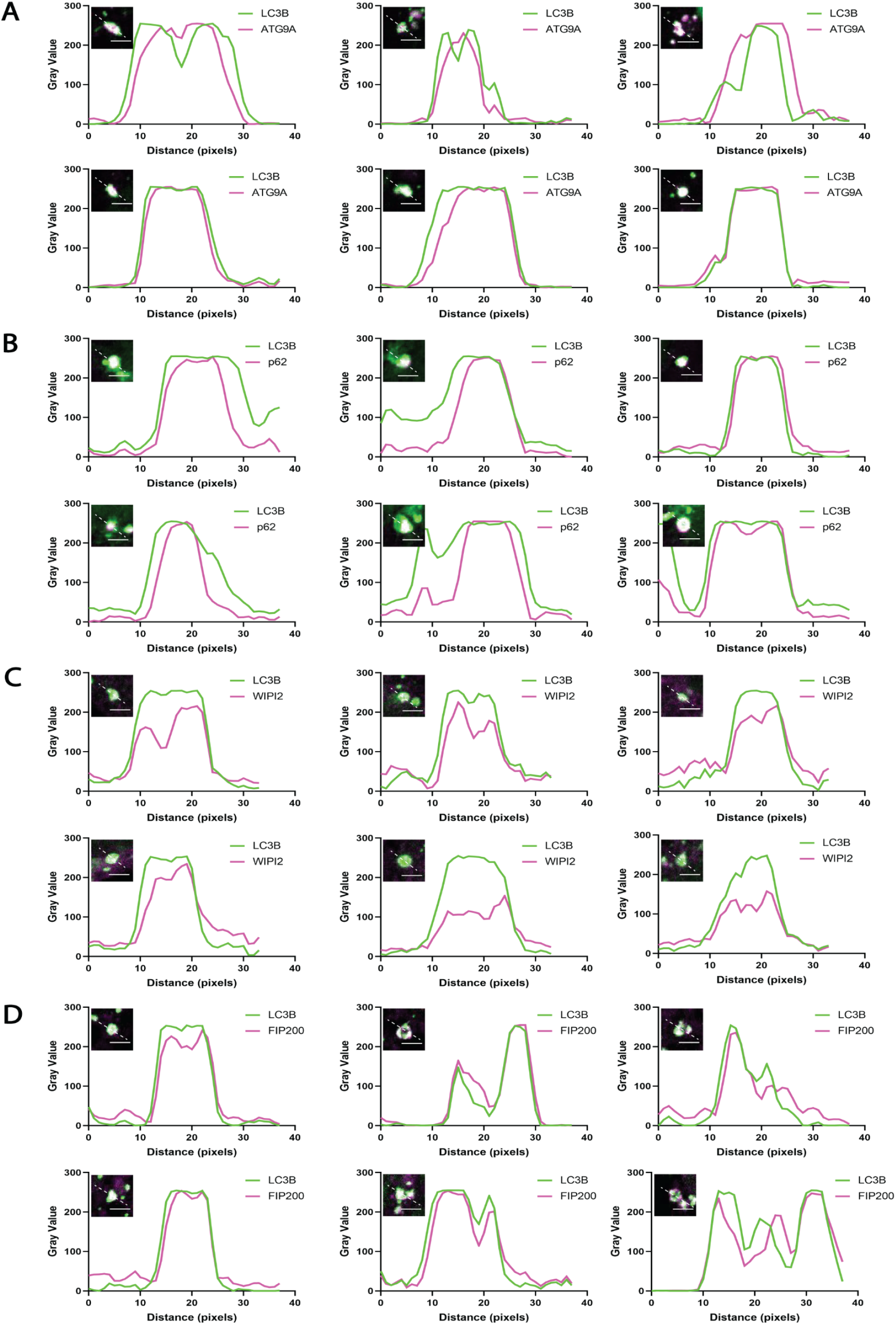
The pre-ATG2 compartment is an organized collection of pre-autophagosomal proteins and membranes. Line scans corresponding to six individual LC3B-positive structures from ATG2 DKO HEK293 cells per early autophagy factor. **(A)** ATG9A signal largely colocalizes with LC3B but in the largest structures is also found enriched in the compartment core. **(B)** p62 consistently colocalizes with LC3B with strongest signal in the center of the compartment in the majority of sampled cells. **(C)** WIPI2 signal is consistently found on the outside of the LC3B signal, colocalizing with LC3B at the rim of the compartment. **(D)** FIP200 colocalizes with LC3B, with both direct overlap and adjacent localization. Inset scale bars: 3 μm.

**Supplemental Figure 2.**
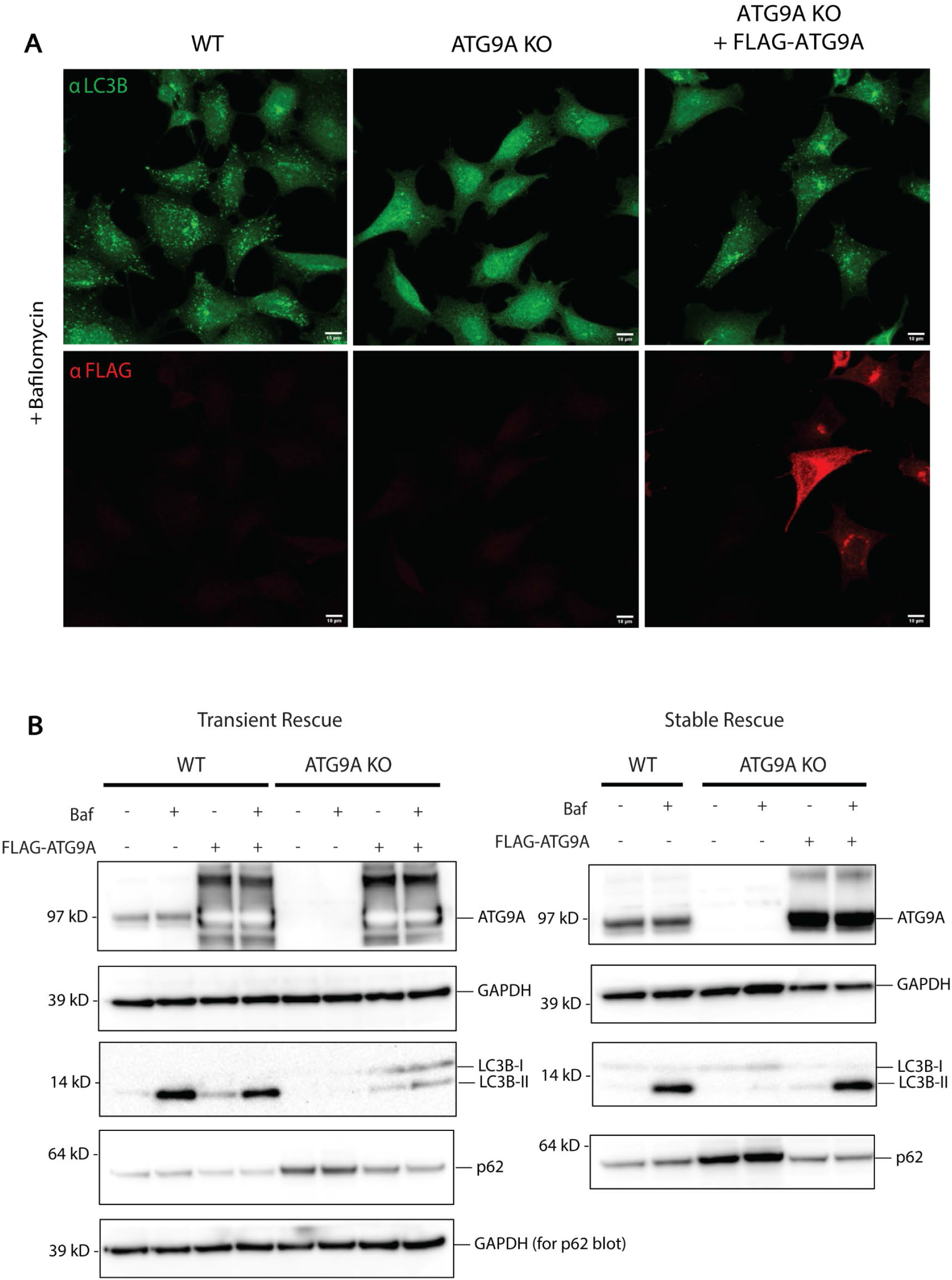
ATG9A knockout displays little to no lipidation. **(A)** Immunofluorescence of WT HEK293 cells compared to ATG9A KO HEK293 cells and ATG9A KO cells rescued with expression of FLAG-ATG9A. Rescue of the ATG9A KO shows restoration of Bafilomycin A1-dependent accumulation of autophagosomes as in the WT. Scale bars: 10 μm. **(B)** Immunoblots showing LC3B lipidation, p62 turnover, and Bafilomycin A1-induced accumulation of lipidated LC3B is restored by addition of exogenous ATG9A in both transient (left) and stable (right) expression in ATG9A KO cells.

**Supplemental Figure 3.**
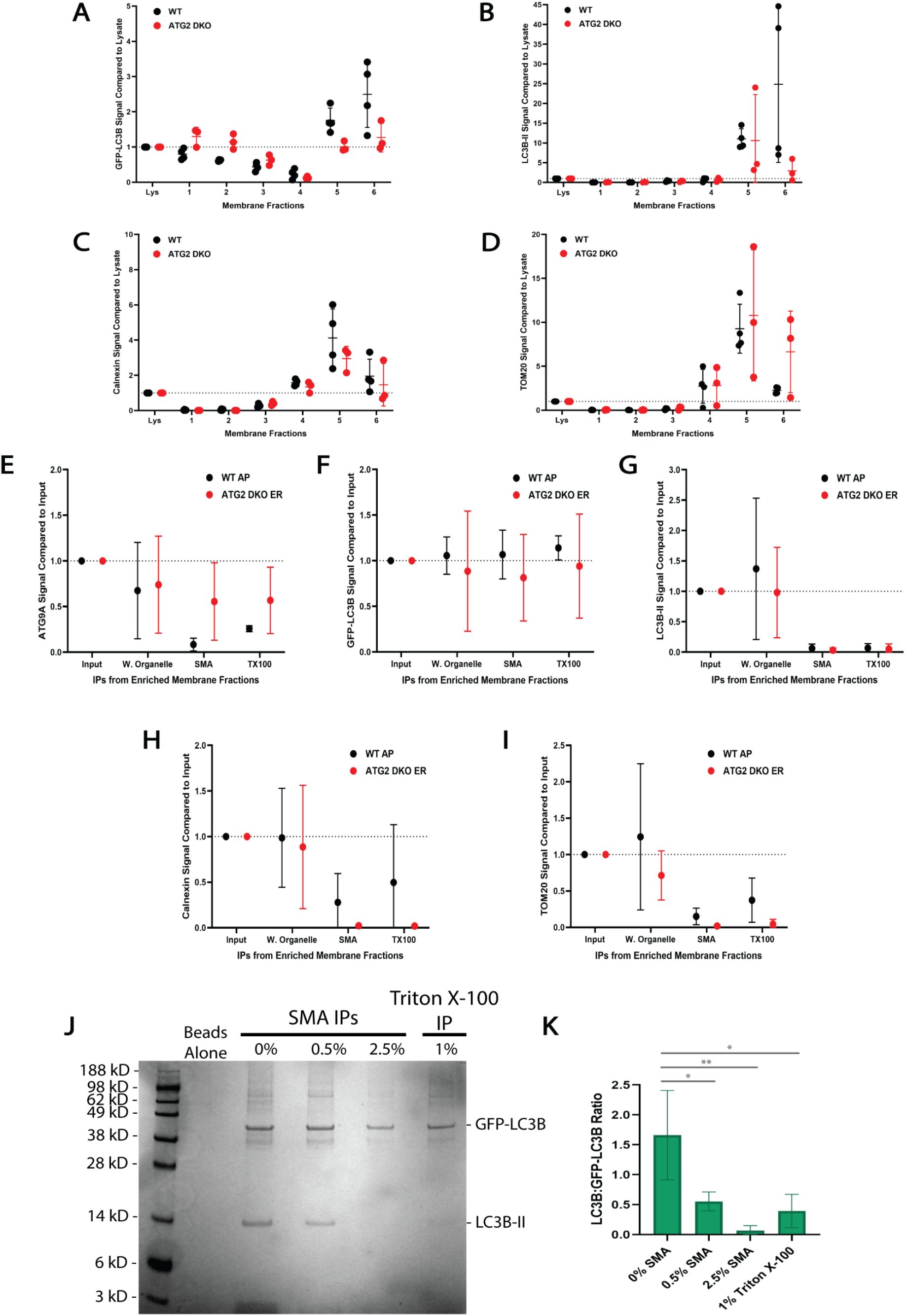
Further quantifications and in vitro controls. **(A-D)** Densitometric quantification of GFP-LC3B, Calnexin, and TOM20 from the density gradient membrane fractions in **Figure 3**. The intensity of the bands were normalized to the lysate, and statistical significance was assessed by multiple unpaired t-tests. *, adjusted p-value < 0.05. Comparisons are nonsignificant unless noted otherwise. **(E-I)** Densitometric quantification of the GFP-LC3B IPs in **Figure 5** comparing protein levels from ATG9A/LC3B-enriched fractions between WT and ATG2 DKO HEK293 cells. The intensity of the bands were normalized to the input, and statistical significance was assessed by multiple unpaired t-tests. All comparisons between WT and ATG2 DKO IPs were nonsignificant. **(J-K)** *In vitro* SMA isolation of GFP-LC3B and LC3B decorated liposomes reveal low frequency of co-residency in SMALPs of randomly distributed proteins. 25 nm liposomes were lipidated as described before (Kauffman et al., 2018), with both LC3B and GFP-LC3B. **(J)** Coomassie-stained SDS-PAGE gel showing IPs of lipidated liposomes treated with various SMA concentrations or Triton X-100. The 2.5% SMA condition mimics that used on the natural membrane SMA IPs in **Figure 5**. The *in vitro* results show that recovery of two distinct peripheral proteins together is relatively rare and further these results closely mirror the recovery of these two proteins from natural sources as shown with the SMA titrations in **Figure 4** on levels of GFP-LC3B and LC3B recovery. **(K)** Densitometric quantification of the LC3B:GFP-LC3B ratio between IPs. Statistical significance was assessed by one-way ANOVA. *, adjusted p-value < 0.05. **, adjusted p-value < 0.01.

**Movie S1. FIB-SEM revealed a spherical compartment of vesicles in ATG2 DKO cells.** FIB-SEM moving down the Z axis of the compartment of vesicles. Scale bar: 500 nm. FIB-SEM slice thickness: 7 nm.

